# Integrative holo-omic data analysis predicts interactions across the host-microbiome axis

**DOI:** 10.1101/2025.10.21.683681

**Authors:** J Merkesvik, J Langa, C Pietroni, A Alberdi, LL Poulsen, AM Bojesen, T Meuronen, S Turunen, O Kärkkäinen, B Westereng, PB Pope, TR Hvidsten

**Affiliations:** Faculty of Chemistry, Biotechnology, and Food Science; Norwegian University of Life Sciences; Ås, Norway; Faculty of Science and Technology; University of the Basque Country; Leioa, Spain; Centre for Evolutionary Hologenomics; University of Copenhagen; Copenhagen, Denmark; Department of Veterinary and Animal Science, University of Copenhagen; Frederiksberg C, Denmark; Afekta Technologies Ltd; Kuopio, Finland; School of Pharmacy; University of Eastern Finland; Kuopio, Finland; Faculty of Biosciences; Norwegian University of Life Sciences; Ås, Norway; Centre for Microbiome Research; Queensland University of Technology; Woolloongabba, Australia

## Abstract

Understanding the interplay between host organisms and their microbiomes is central to the development of sustainable food systems. However, high dimensionality and spurious associations remain major obstacles to extracting meaningful biological insight from multi-omic host-associated microbiome data; a challenge further exacerbated when “holo-omic” analyses across the host-microbiome boundary is considered. Here, we show that a computational method designed for multi-omic analysis in eukaryotes can be leveraged to integrate and analyse five layers of holo-omic data from porcine hosts and their gut microbiomes. We collected caecal tissue and digesta samples during a feeding trial that tested the impact of microbiota-directed fibres (acetylated galactoglucomannan) at critical developmental stages. From 800,000 features including microbial and host genes, metagenome-assembled genomes, and metabolites from caecal tissue and digesta, we used multiset correlation and factor analysis to select the most relevant features for capturing coordinated patterns across omic layers. From these features, we predicted over 2,000 putative host-microbiome interactions based on co-occurrence. Some of them reflected previously known relationships between animal and microbiome features, such as microbial genes for carbohydrate metabolism being linked to glycoside abundances in host tissue. Other predicted co-occurrences included features that were not detected in single-omic analysis and offer new hypotheses of host-microbiome interactions that warrant future investigation. Hence, we showcase an application of holo-omic analysis that avoids common pitfalls in high-dimensional data analysis; identifies known interactions as a form of validation; and most importantly, predicts new leads for understanding host-microbiome symbiosis.

**Importance:** While study systems involving mammalian hosts and their microbiomes are inherently complex, multi- and holo-omic analyses promise to provide interpretable results with translational value for the animal production industry. Unfortunately, computational methods capable of this kind of integration are currently scarce, as most existing multi-omics approaches have been developed for analysis of data layers within a single multicellular organism. We propose to adapt existing multi-omic methods for holo-omics by combining feature selection and interaction inference. This two-step analysis approach addresses common challenges in data-driven studies and can be implemented with a variety of tools for feature selection and interaction modelling. Through this holistic approach, we show that both known and novel relationships across the holobiont axis can be identified in a data-driven manner, offering new targets for the continued study of host-microbiome interactions and the effect of dietary interventions on production animals.

## Observations

Multi-omic studies investigating biomolecules from cells and tissues have over the past decade provided important insight into the organisation and regulation of complex biological systems. Holo-omics is an extension of multi-omics where a host organism and its associated microbiomes are represented by separate data layers and co-analysed [1]. Although capturing perspectives from both the host and its microbiomes is essential for studying cross-kingdom interactions, holo-omic datasets present several challenges for researchers. High-dimensional feature spaces are often computationally expensive and overwhelming to analyse, interpret, and visualise; the addition of features rarely entails notable increases in sample size to prevent overfitting; and data heterogeneity calls for omic-specific pre-processing to ensure the validity of inferred results [1,2]. Moreover, coincidental associations must be distinguished from causal relationships to identify biological processes underlying host-microbiome interactions [2]. Several tools have been developed for multi-omics (a community-assembled list being available on https://github.com/mikelove/awesome-multi-omics), but a gold-standard methodology has yet to be established [1]. Even fewer methods have been tailored to predict interactions across the host-microbiome boundary.

We sought to observe the potential of holo-omic methodologies to better predict how complex microbiomes interact with their host and used a defined porcine nutrition experiment as a test system (**Fig. 1**). Pigs are an increasingly important livestock [3], but great changes in gut physiology during weaning make piglets vulnerable to potentially fatal gastrointestinal diseases [4–6]. As the gut microbiome plays an integral role in host health [7–10], the development of technologies to stimulate beneficial gastrointestinal ecosystems can address several industry challenges. Members of the endogenous gut microbiota in mature pigs can ferment dietary fibres into host-available compounds, such as beneficial short-chain fatty acids (SCFAs) [4– 7,9]. In this context, microbiota-directed fibres (MDFs) can be administered to target and selectively promote microbial commensals in the gut that improve the health and well-being of the host animal during weaning [4–7]. MDF administration has been showcased using the soft-wood hemicellulose acetylated galactoglucomannan (AcGGM) (**Fig. 1A**), which targets SCFA-producing commensals like *Bacteroides, Bifidobacterium, Blautia, Butyricicoccus, Butyrivibrio, Faecalibacterium, Prevotella, Roseburia*, and *Ruminococcus* [4,6–8,12,14–16]. Select members of these genera harbour AcGGM-specific polysaccharide utilisation loci (PULs) with carbohydrate-active enzymes (CAZymes) that elicit AcGGM degradation, including carbohydrate esterases (CEs) and glycoside hydrolases (GHs). Despite the strong modulative effects of AcGGM in the porcine gut microbiome, its system-wide impact beyond the target species is poorly understood and thus justified an alternative holo-omic investigative approach (**Fig. 1D**). Therefore, we generated metagenomics, meta- and host transcriptomics, and untargeted metabolomics from both microbiome and host tissue samples to study the impact of AcGGM introduced to the pig host and its gut microbiome at different developmental stages.

**Figure 1.**
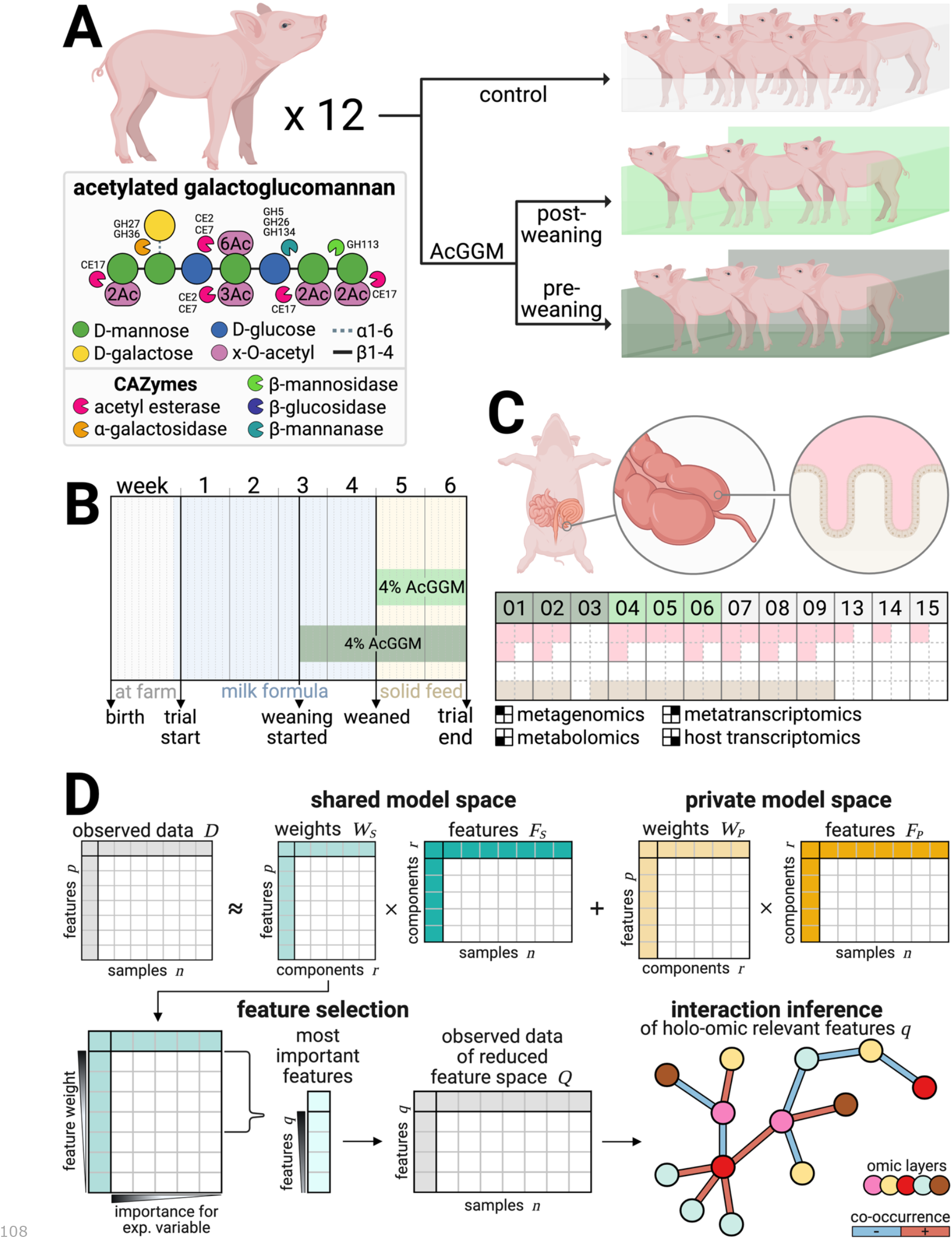
Experimental design, data generation, and holo-omic analysis procedure of the porcine feeding trial with AcGGM. **A)** Division of 12 piglets into three groups of individually housed piglets: one control, and two groups introduced to AcGGM at different developmental stages. **B)** Trial timeline reporting milestone events and porcine diets. Solid feed composition is found in Suppl. 1. **C)** Sites for sample collection after 6 weeks – caecal digesta (pink) and gut wall (tan) – and omic data layers generated from these samples. Sampling and data generation steps are described in Suppl. 2. **D)** Proposed two-step methodological framework for predicting host-microbiome interactions. Using multiset correlation and factor analysis (MCFA) [11], observed data (D) are reconstructed as the sum of a shared space (W_S_ × F_S_) capturing common data patterns, and private model spaces for each omic layer (W_P_ × F_P_) (adapted from [1]). The fitted weights in the shared space (W_S_) are then used to select features with the most relevance for capturing cross-omic patterns in the dataset. The observed omic data of the reduced feature space (Q) is used to predict direct interactions between host- and microbiota-associated features. Abbreviations: AcGGM: acetylated galactoglucomannan, CAZymes: carbohydrate-active enzymes. Figure created in BioRender: Merkesvik J. (2025) https://biorender.com/e0obj4f.

Analysis of the porcine study system at the single-omic level showed that the caecal microbiota appeared matured in all piglets irrespective of AcGGM administration. Typical genera for the infant gut – *Lactobacillus, Bacteroides, Clostridium*, and *Escherichia* – had low relative abundances (0.9%, 0.3%, 0.3%, and 0.3%, respectively) compared to taxa associated with mature animals: *Prevotella, Alloprevotella, Agathobacter, Blautia, Faecalibacterium,* and *Roseburia* (6.2%, 2.8%, 2.6%, 1.4%, 1.4%, and 1.3%, respectively) [4,6,7]. Still, we observed significant differences (false discovery rate-adjusted (FDR) p-value<0.05) in piglets introduced to AcGGM at different developmental stages. Compared to control and post-weaning AcGGM animals, pre-weaning AcGGM microbiomes displayed higher abundances of *Ruminiclostridium sp.* and *Basfia porcinus* (absolute log2 fold change (LFC)>6), *Lactobacillus johnsonii* (LFC>5), *Butyribacter sp.* and *Alloprevotella sp.* (LFC>4), *Prevotella pectinovora* (LFC>3), and *Duodenibacillus sp.* (LFC>2) (**Suppl. 3**). The transcriptomic activity of these taxa mirrored their abundance increases, suggesting that early AcGGM introduction influenced both the composition and functional activity of the caecal microbiota. A selection of host-associated metabolites and genes also displayed differential abundance and expression across diet groups. These features could however not straight-forwardly be connected to observations from the microbiota, preventing us from inferring host-microbiome interactions relevant for AcGGM supplementation.

To predict biologically relevant interactions across the host-microbiome axis, we implemented a two-step analysis approach. First, we used multiset correlation and factor analysis (MCFA) [11] for feature selection. The tool jointly modelled all five observed omic layers, formulating so-called “shared” components (*F_S_*) which – when scaled by feature-specific weights (*W_S_*) – capture data patterns common across input layers (**Fig. 1D**). Subsequently, omic-specific “private” components (*F_P_*) and weights (*W_P_*) were fit to explain the remaining variance in each layer. The one-dimensional shared space captured 9-29% of observed variance across the input layers (**Fig. 2A**) and aligned well with the developmental stage for AcGGM introduction (pseudo-R^2^>0.6) (**Fig. 2B**). We therefore used the shared feature weights (*W_S_*) as indicators of importance for coordinated data patterns. By extracting features with absolute shared weight in the 93^rd^ percentile from each layer, we narrowed down the feature space from 806,854 to 56,442 features. These were considered the most relevant features for describing cross-omic data patterns in response to AcGGM introduction.

**Figure 2.**
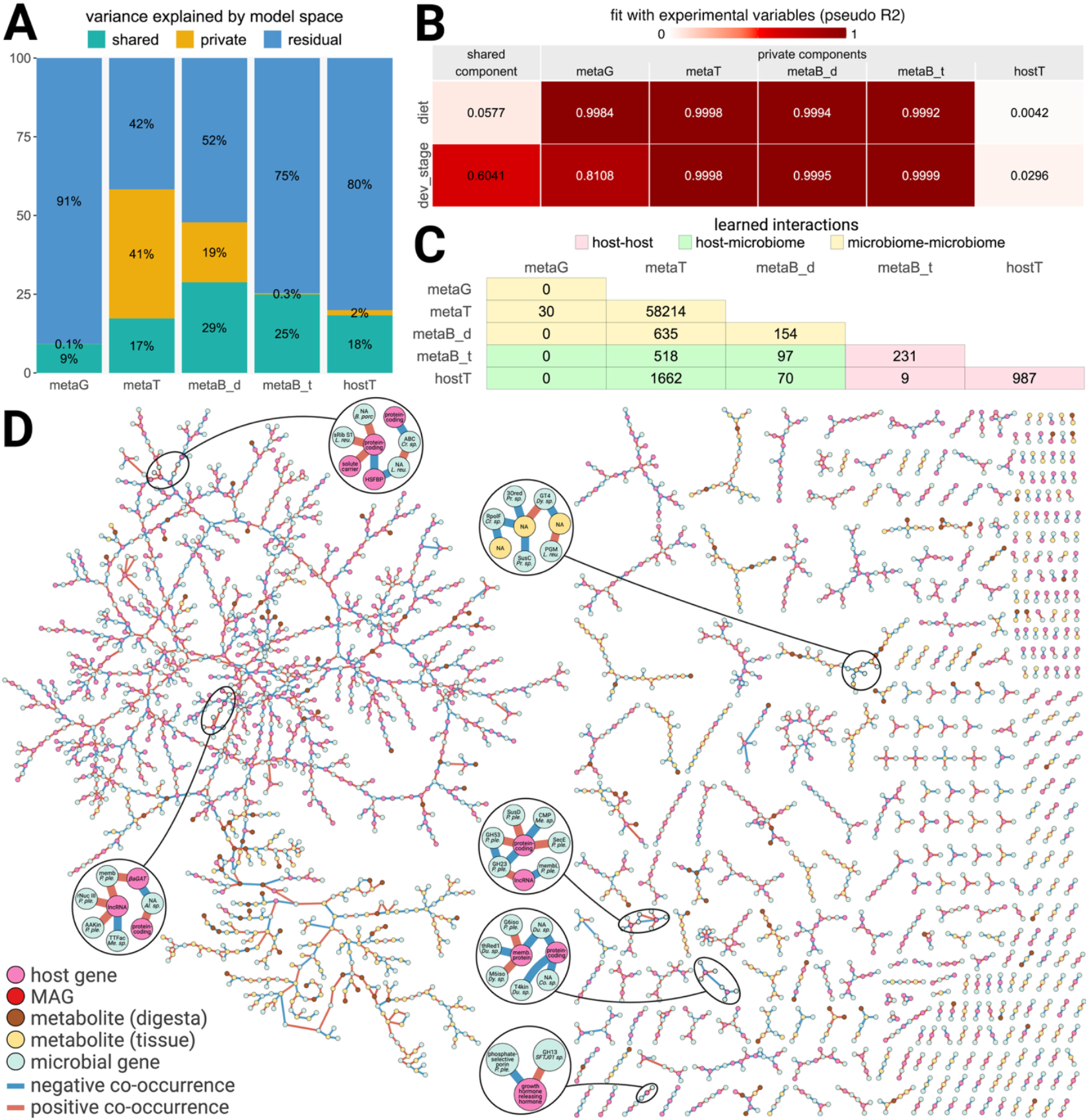
Outcomes of holo-omic integration through feature selection and direct interaction inference. **A)** Percent variance explained by model spaces for each input omic data layer. Style adapted from [11]. **B)** Ratio of sample metadata variance explained by each learned model space, calculated as the improvement in log likelihood from a logistic regression null model to a fitted model. **C)** Number of predicted interactions between the most important features for cross-omic data patterns, limited to the main component of the inferred co-occurrence network. Cell colour classifies the interaction as specific to host (pink) or microbiome (yellow) omic layers, or as crossing the holobiont axis (green). **D)** Subset of co-occurrence network of omic features with most impact on shared model space, only including nodes with host-microbiome interactions in the main connected component (green cells in panel C). Node colour represents omic layer and edge colour conveys direction of co-occurrence. Selected predicted interactions are highlighted in subpanels. The full graph is available on GitHub. Abbreviations: metaG: metagenomics, metaT: metatranscriptomics, metaB_d: metabolomics digesta, metaB_t: metabolomics tissue, hostT: host transcriptomics, dev_stage: development stage, 3Ored: 3-oxoacyl-(acyl-carrier protein) reductase, AAKin: amino acid kinase, ABC: adenosine triphosphate-binding cassette, Al: Alloprevotella, βaGAT: beta-1,4-N-acetyl-galactosaminyltransferase, B. porc: Butyricicoccus porcorum, CMP: 3-deoxy-manno-octulosonate cytidylyltransferase, Co: Copromonas, Cr: Cryptobacteroides, Du: Duodenibacillus, Dy: Dysosmobacter, G6iso: glucose-6-phosphate isomerase, GH: glycoside hydrolase, GT: glycosyl transferase, HSFBP: heat shock factor binding protein, L. reu: Limosilactobacillus reuteri, lnc: long non-coding, M6iso: mannose-6-phosphate isomerase, Me: Mediterranea, memb: outer membrane protein, membL: outer membrane lipoprotein, PGM: phosphoglucomutase, P. ple: Phocaeicola plebius, Pr: Prevotella, rNuc: ribonuclease, RpolF: RNA polymerase sigma-70 factor, SecE: protein translocase subunit, sRib: small subunit ribosomal protein, Sus: starch utilisation system, T4kin: tetra-acyl-disaccharide 4’-kinase, thRed: thioredoxin, TTFac: transcription termination factor. Figure assembled in BioRender: Merkesvik J. (2025) https://biorender.com/z5g5s31.

Next, we used the selected omic features as input to FlashWeave [17] to predict interactions through co-occurrence. Variables on diet and developmental stage were included as covariates to eliminate indirect associations driven by experimental and technical factors. We identified 70,888 putative relationships between these features, of which 47,327 nodes and 62,609 edges were contained in one connected network component. Here, 94% of predicted interactions occurred between microbiome features; 2% connected host features; and 4% crossed the host-microbiome boundary (**Fig. 2C**). Limiting the graph to features with host-microbiome interactions yielded a subnetwork with 3,276 nodes and 3,011 links (**Fig. 2D**). The majority of these putative interactions (72%) connected microbial and host genes, and 10% were between microbial genes and host-associated metabolites. Here we observed enrichments of genes from *Faecivivens sp., Limosilactobacillus reuteri*, and *Phocaeicola plebeius.* Several CAZymes were among these features – including nine GHs; a starch utilisation system D (SusD) carbohydrate transporter in *P. plebeius*; and phosphoglucomutase in *F. sp*. - suggesting that carbohydrate degradation was relevant for host-microbiome interactions in our system. Moreover, several predicted host-microbiome interactions involved features from metagenome-assembled genomes (MAGs) that were more abundant in the pre-weaning AcGGM animals, as identified by single-omic analysis. For instance, the expression of 51 *Duodenibacillus sp.* genes were linked to 28 tissue metabolites (e.g., the glycoside sachaloside, isoferulic acid, and linoleyl carnitine) and 33 host genes (including membrane and binding proteins); while a dehydrogenase from *Lactobacillus johnsonii* negatively co-occurred with macrolide/pyrrolizine in host tissues. Our holo-omic approach thus recapitulated features flagged by single-omic analyses and provided additional features and hypothesised interactions of interest for the continued study of the effect of AcGGM on porcine holobionts. The data-driven feature selection and interaction inference demonstrated here may prove valuable for other high-dimensional heterogeneous multi-omic studies and underscore the need for holistic approaches when analysing complex biological systems.

In summary, we applied software developed for multi-omics on a holo-omic dataset to predict interactions between host- and microbiome-associated features with implied importance for coordinated data patterns in response to AcGGM administration. First, feature selection with MCFA circumvented issues of high-dimensionality and data heterogeneity. Secondly, FlashWeave predicted interactions between selected features while disregarding spurious associations driven by experimental and technical factors. The resulting co-occurrence network not only reiterated findings from our single-omic analyses but also suggested novel features and hypothetical interactions to consider for studies that validate the effect of AcGGM on pre-weaning piglets.

## Abbreviations

AcGGM: acetylated galactoglucomannan
CAZyme: carbohydrate-active enzyme
CE: carbohydrate esterase
FDR: false discovery rate
GH: glycoside hydrolase
LFC: log2 fold change
MAG: metagenome-assembled genome
MCFA: multiset correlation and factor analysis
MDF: microbiota-directed fibre
PUL: polysaccharide utilisation locus
SCFA: short-chain fatty acid
Sus: starch utilisation system

## Acknowledgements

We acknowledge the financial support of the European Union’s Horizon 2020 research and innovation programme under the grant agreement 101000309-3D’omics. JL was funded by the Postdoctoral Program for the Improvement of Doctoral Research Staff of the Basque Government (grant number POS_2022_1_0011). We thank the technicians and PhD candidates of the BioRef group (Norwegian University of Life Science, Norway) for AcGGM production; personnel at the Department of Veterinary and Animal Science (University of Copenhagen, Denmark) for conducting the animal trial; and lab technicians and bioinformaticians at the Centre for Evolutionary Hologenomics (UCPH, Denmark) for sample preparation and data generation. Lastly, we acknowledge the Orion High Performance Computing Centre at the Norwegian University of Life Sciences for providing computational resources that have contributed to the research results reported within this paper.

## Data availability

Raw sequencing data from metagenomics, transcriptomics, and metatranscriptomics are available in the European Nucleotide Archive under accession PRJEB100724, as part of the umbrella project 3D’omics (PRJEB86267). Raw metabolomics data are found on MetaboLights under accession REQ20251014213828. Count tables, metadata, scripts, network files, and an RMarkdown report detailing the analysis procedure are found on GitHub (jennymerkesvik/3domics_wp3-2). Supplementary materials include detailed method descriptions and additional data visualisations used herein to report single-omic analysis results.

