## supplementary_materials for "Integrative holo-omic data analysis predicts interactions across the host-microbiome axis"

S3.1: Trial metadata on animal performance and feed administration
S3.2: Phylogenetic tree of MAG catalogue
S3.3: Differentially abundant MAGs across diet groups
S3.4: Differentially abundant MAGs across developmental stages
S3.5: Differentially abundant metabolites across developmental stages
S3.6: Differentially expressed host genes across diet groups
S3.7: Differentially expressed host genes across developmental stages
References........ 21

#### Abbreviations

| | |
|---|---|
| AcGGM | acetylated galactoglucomannan |
| FDR | false discovery rate |
| HPLC | high-performance liquid chromatography |
| LC-MS | liquid chromatography mass spectrometry |
| LFC | log2 fold change |
| MAG | metagenome-assembled genome |
| MCFA | multiset correlation and factor analysis |
| MS/MS | tandem mass spectrometry |
| PCR | polymerase chain reaction |
| rD-ratio | robust dispersion ratio |
| rRSD | robust relative standard devioation |
| UHPLC | ultrahigh-performance liquid chromatography |

#### Suppl. 1: Porcine solid feed composition

Composition of the AgroSoft solid feed given to all animals during the trial.

| nutrient | unit | per kg | per energy | nutrient | unit | per kg | per energy |
|---|---|---|---|---|---|---|---|
| FEsvin ny | FEsv | 1.219 | 1.0 | potassium | g | 7.570 | 6.210 |
| FE so ny | FEso | 1.199 | 0.983 | chloride | g | 6.370 | 5.220 |
| crude protein | % | 18.280 | 14.990 | vitamin A, additive | 1000 i.e | 16.0 | 13.120 |
| nitrogen | % | 2.930 | 2.40 | vitamin D3, additive | 1000 i.e | 2.0 | 1.640 |
| lignin | % | 2.870 | 2.350 | vitamin E (synthetic), additive | i.e | 100.0 | 82.0 |
| crude fat | % | 6.650 | 5.450 | vitamin E/DL alpha tocopherol, additive | mg | 91.0 | 74.620 |
| crude ash | % | 5.310 | 4.350 | vitamin E eq. (from Proviox) | mg | 100.0 | 82.0 |
| dry matter | % | 88.670 | 88.670 | vitamin K3, additive | mg | 2.0 | 1.640 |
| starch | g | 367.020 | 300.960 | vitamin B1/thiamine, additive | mg | 10.0 | 8.20 |
| sugar | g | 63.40 | 51.990 | vitamin B2/riboflavin, additive | mg | 5.0 | 4.10 |
| st.dig. crude protein | g | 166.630 | 136.640 | vitamin B6/pyridoxin, additive | mg | 5.0 | 4.10 |
| st.dig. lysine | g | 13.170 | 10.80 | vitamin B12, additive | mg | 0.050 | 0.040 |
| st.dig. methionine | g | 5.160 | 4.230 | D-pantothenic acid, additive | mg | 35.0 | 28.70 |
| st.dig. methionine+cystine | g | 7.770 | 6.370 | niacin, additive | mg | 25.0 | 20.50 |
| st.dig. threonine | g | 9.030 | 7.410 | biotin vitamin H, additive | mg | 0.30 | 0.250 |
| st.dig. tryptophan | g | 2.770 | 2.270 | choline chloride, additive | mg | 350.0 | 287.0 |
| st.dig. isoleucine | g | 6.530 | 5.350 | folinic acid, additive | mg | 3.0 | 2.460 |
| st.dig. valine | g | 8.670 | 7.110 | L-carnitine | mg | 23.0 | 18.860 |
| lysine | g | 13.970 | 11.450 | betaine hydrochloride, additive | mg | 100.19 | 82.160 |
| methionine | g | 5.380 | 4.410 | Fe, iron II sulphate | mg | 120.0 | 98.40 |
| threonine | g | 9.820 | 8.050 | Cu, copper II sulphate | mg | 130.0 | 106.60 |
| tryptophan | g | 2.990 | 2.450 | Mn, manganese oxide | mg | 70.0 | 57.40 |
| calcium | g | 6.460 | 5.30 | Zn, zink sulphate | mg | 85.0 | 69.70 |
| calcium formate | mg | 0.0 | 0.0 | I, calcium iodate | mg | 3.06 | 2.510 |
| phosphorus | g | 6.720 | 5.510 | Se, sodium selenite | mg | 0.25 | 0.210 |
| dig. phospgorus, 0% phytase | g | 3.820 | 3.130 | 6-phytase (EC 3.1.3.26) | OTU | 265.0 | 217.30 |
| dig. phospgorus, 60% phytase | g | 4.080 | 3.350 | endo-1,4-beta-xylanase | EPU | 1500.0 | 1230.020 |
| dig. phospgorus, 100% phytase | g | 4.190 | 3.430 | beta glucanase (EC 3.2.1.5) | 1000U | 0.10 | 0.080 |
| dig. phospgorus, 150% phytase | g | 2.280 | 3.510 | BHT (antioxidant) | mg | 10.0 | 8.20 |
| dig. phospgorus, 200% phytase | g | 4.350 | 3.560 | antioxidant | mg | 21.52 | 17.650 |
| sodium | g | 3.060 | 2.510 | propyle gallate | mg | 5.0 | 4.10 |
| magnesium | g | 1.450 | 1.190 | ProHacid® | mg | 7000.0 | 5740.080 |
| magnesium sulphate | g | 0.0 | 0.0 | | | | |

### Suppl. 2: Method descriptions

#### Acetylated galactoglucomannan fibres

AcGGM was produced following the protocol described in Michalak *et al*. [1]. In summary, dry wood from Norway spruce (*Picea abies*) was milled into chips and soaked in a solution with sodium citrate and potassium phosphate buffers. The wood chips were steam exploded at 200°C and 14.5 bar in a 20 L pressure vessel, using a 25 kW electric boiler (Parat, Norway) for steam production. Water was then added to create a slurry, which was filtered to separate the water-insoluble material from the wood chips. The solid mass was dried at 100°C for 48 hours and cut into smaller pieces using a cutting mill (Retsch, Germany) with a 0.5 mm sieve.

#### Animal trial and sampling

The feeding trial was conducted at the University of Copenhagen during spring 2022 under the experimental animal license number 2020-15-0201-00520. Twelve ten-day old male piglets (2.8-4.9 kg) of the same litter from Bøgholt farm (Blangstrupsvej 17, DK-5610 Assens) were housed individually and divided into three weight-balanced experimental groups with specific feeding regimens over a period of six weeks. All received the same basal diet: four weeks of DanMilk Supreme 1.0 milk replacement formula (AB Neo, Denmark), and two weeks of solid feed (AgroSoft, Denmark) (**Suppl. 1**). All piglets were weaned at age 24 days. Two of three groups received AcGGM fibres as a supplement at 4% inclusion, but starting at different time points: one for the final four weeks of the trial, spanning two weeks before and two weeks after weaning; the second only for the final two trial weeks, starting after weaning. Body weight, feed intake, and clinical scores are visualised in **Fig. S3.1**. After the trial's sixth week, the piglets were euthanized by Zoletil anaesthesia (0.1 mL/kg) followed by a lethal intracardial pentobarbital dose (0.25 mL/kg).

#### Sample collection and preparation

Endpoint samples were collected from the caecum of each animal. Digesta (100 mg) was sampled in all 12 animals for metagenomic, metatranscriptomic, and untargeted metabolomics analyses, whereas tissue samples (1x1 cm) were collected from nine individuals for transcriptomic and untargeted metabolomics analyses. Samples intended for metagenomics and (meta)transcriptomics were stabilised in DNA/RNA Shield (1 mL) (Zymo, USA) to prevent nucleic acid degradation and enable long-term storage at -20°C until further processing for data generation. For untargeted metabolomics, digesta and tissue samples were snap frozen in liquid nitrogen and stored at -80°C until further processing.

Samples for metagenomics, metatranscriptomics, and host transcriptomics were prepared following the laboratory workflow of the Earth Hologenome Initiative (https://www.earthhologenome.org/laboratory). Briefly, thawed samples were mechanically lysed using a Lysing Matrix E (MP Biomedicals, USA) with two 6-minute bead-beating cycles at 30 Hz on a TissueLyser II (Qiagen, Germany). Silica magnetic beads and solid-phase reversible immobilisation were used to isolate RNA and DNA in separate fractions and to remove inhibitors. DNA and RNA concentrations were quantified with a Quibit 3 Fluorometer using Quibit HS or BR Assay Kits (Thermo Fisher Scientific, USA). For metagenomics, DNA was fragmented to ~400 bp by ultrasonication using a Covaris LE220 platform. Library preparation was conducted following a blunt-end adapter ligation single-tube protocol [2] with optimised conditions as described by Mak *et al.* [3]. qPCR screening was used to determine the number of PCR cycles needed for library indexing. Samples were then amplified using unique dual index primers before the libraries were analysed through capillary electrophoresis with Fragment Analyzer (Agilent, USA). Metagenomic libraries were generated using the Illumina Stranded Total RNA Prep kit (Illumina, USA) with enzymatic rRNA depletion via Ribo-Zero Plus (Illumina, USA). Transcriptomic libraries employed bead-based rRNA depletion using the TruSeq Stranded Total RNA Library Prep kit for Human/Mouse/Rat (Illumina, USA). Sequencing libraries were pooled and sequenced with an Illumina NovaSeq6000 system using S4 flow cells, generating approx. 10 GB per (meta)transcriptomic library, and 5 GB per metagenomic library.

For untargeted metabolomics, digesta and tissue samples were transferred to cooled Bead Ruptor homogeniser tubes (Omni International, USA) with four metal beads and cold methanol (80%, 400 µL/100 mg tissue, 1000 µL/100 g digesta). Digesta samples were homogenised using a Bead Ruptor 24 Elite (Omni International, USA) (2±2°C, 7.0 m/s, 2x30 seconds, 20 seconds between cycles). Tissue samples followed the same homogenisation protocol repeated once. Samples were incubated on ice for 15 minutes before being vortexed (10 seconds) and centrifuged (4°C, 17,000 G, 10 minutes). Supernatants were decanted through a 0.2 µm Acrodisc filter with polytetrafluoroethylene membrane (Pall, USA) into HPLC vials (Agilent, USA) for LC-MS analysis. Quality control samples – one for digesta and one for tissue samples – were prepared by pooling 150 µL of supernatants from each sample of each respective sample type into another HPLC vial. The two pooled samples were vortexed and filtered like the other samples. LC-MS analysis utilised a 1290 Infinity II UHPLC (Agilent, USA) coupled with a high-resolution quadrupole time of flight Agilent 6546 mass spectrometer with a Dual Jet Stream ion source (Agilent, USA) and followed the protocol described in Hanheniva *et al.* [4] and Klåvus *et al*. [5]. In brief, a Zorbax Eclipse XDB-C18 column (2.1×100 mm, 1.8 µm) (Agilent, USA) was used for reversed-phase separation, while hydrophilic interaction liquid chromatography separation was conducted with an UHPLC ethylene bridged hybrid amide column (Waters, USA). After each chromatographic run, samples were ionised through Jet Stream electrospray in positive and negative mode, yielding four data files per sample. Collision energies for MS/MS analysis were selected as 10, 20, and 40 V for compatibility with spectral databases.

#### Omic data generation

Metagenomics, metatranscriptomics, and host transcriptomics sequencing data were processed using Snakemake [6] workflows developed within the 3D'omics project, which are available on GitHub (https://github.com/3d-omics).

For untargeted metabolite data generation, peak detection and alignment was performed in MS-DIAL (ver. 4.92) [7]. Mass-to-charge ratios between 50 and 1,500 at any retention time were considered for peak collection. Minimum peak height amplitude was set to 2,000, and detection used the linear weighted moving average algorithm. To align peaks across samples, tolerances were set to 0.2 min retention time and 0.015 Da m/z. Solvent background was removed using solvent blank samples with the condition that the maximum signal abundance across samples was at least five times higher than the mean signal in the blanks. A total of 62,526 metabolite features remained after peak picking, and were subjected to further preprocessing and clean-up with the R-package notame (ver. 0.2.0) [5] for each sample type separately. Metabolomic features were considered high quality based on presence (>70% of quality control samples and >50% of samples in at least one study group); rRSD (<20%); and rD-ratio (<10%). In addition, features with too high rRSD or rD-ratio were still considered good quality if their classic RSD, rRSD, and basic D-ratio were all low (<10%). Low-quality features were flagged and discarded, leaving 16,328 metabolite features in the dataset.

Compound identification was conducted with MS-DIAL by comparing chromatographic and spectrometric characteristics with both in-house and publicly available databases (MassBank, ReSpect, RIKEN, GNPS, Fiehn libraries, CASMI2016, MetaboBASE, BMDMS-NP, and PFAS). Additional identifiers were obtained using MS-FINDER (ver. 3.60) [8] to compare acquired MS/MS spectra with *in silico*-generated records for known compounds. All metabolite annotations followed the Metabolomics Standards Initiative by the Chemical Analysis Working Group [9].

The metagenome count matrix was filtered for genome coverage (>50% in at least two samples) and presence in more than one sample, reducing the MAG catalogue from 439 to 373 populations, spanning 815,027 microbial genes. The host genome saw 27,376 mapped genes within these samples, of which 22,170 were expressed in at least two samples and thus included in the analysis. From the 16,284 high-quality metabolic features, 6,388 with acquired MS/MS data were kept in the metabolomic dataset before initiating omic data analyses.

#### Omic data analyses

All statistical analyses were conducted in R (ver. 4.4.2) [10] on an aarch64 Apple Darwin20 platform running macOS Sequoia 15.7, unless stated otherwise. Relevant data and metadata files are available in the GitHub repository jennymerkesvik/3domics_wp3-2, which also contains an RMarkdown report demonstrating the full analysis approach.

Read count-based omic datasets (metagenomics, metatranscriptomics, and host transcriptomics) were compared across sample groups with DESeq2 (ver. 1.46.0) [11], using default settings and significance thresholds of absolute LFC > 1; FDR p-value < 0.05; and base mean > 50. Molecular features from metabolomics were compared across pairs of sample groups using two-tailed t-tests assuming equal variance, false discovery rate-corrected p-values, and log2 fold changes. Variance-stabilising transformations were used to normalise the read count data before they were used for holo-omic integration. Metabolomic features were log2-transformed for the same purpose. R packages ggplot2 (ver. 3.5.1) [12], ggtree (ver. 3.14.0) [13], and ggtreeExtra (ver. 1.16.0) [14] were used to visualise analysis results.

Using Python (ver. 3.12.9) [15], we used MCFA (ver. 1.0.2) [16] to model the holo-omic dataset. The utilised script is found on GitHub, in which standardised omic data layers across common samples were used as input for the MCFA function `fit()`, while querying a one-dimensional shared space (`d=1`). Shared and private components and weights, along with files informing on variance explained by each model space, was imported into R to inspect and visualise model performance. Features with weight for the shared model component in the 93rd percentile were selected as the most relevant features for identifying host-microbiome interactions, while yielding a manageable number of features for predicting cross-omic relationships. A co-occurrence network of these omic data were created with FlashWeave (ver. 0.19.2) [17], which was implemented in Julia (ver. 1.11.6) [18]. Specifically, the function `learn_network()` was provided a matrix consisting of the selected omic data and one-hot encoded metadata on diet group and developmental stage for dietary fibre introduction; a mask to inform on which features were experimental covariates; and additional settings `sensitive = true`, `heterogenous = false`, `n_obs_min = 7`, `normalize = false`. The learned network was imported into Cytoscape (ver. 3.10.3) [19] and R for inspection and visualisation.

#### Suppl. 3: Additional visualisations

Metadata on animal performance and feed administration (**Fig. S3.1**). MAG catalogue composition and differential transcriptomic activity (**Fig. S3.2**). Differentially abundant features across metagenomic (**Figs. S3.3-2**), metabolomic (**Fig. S3.5**), and host transcriptomic (**Figs. S3.6-5**) data layers.

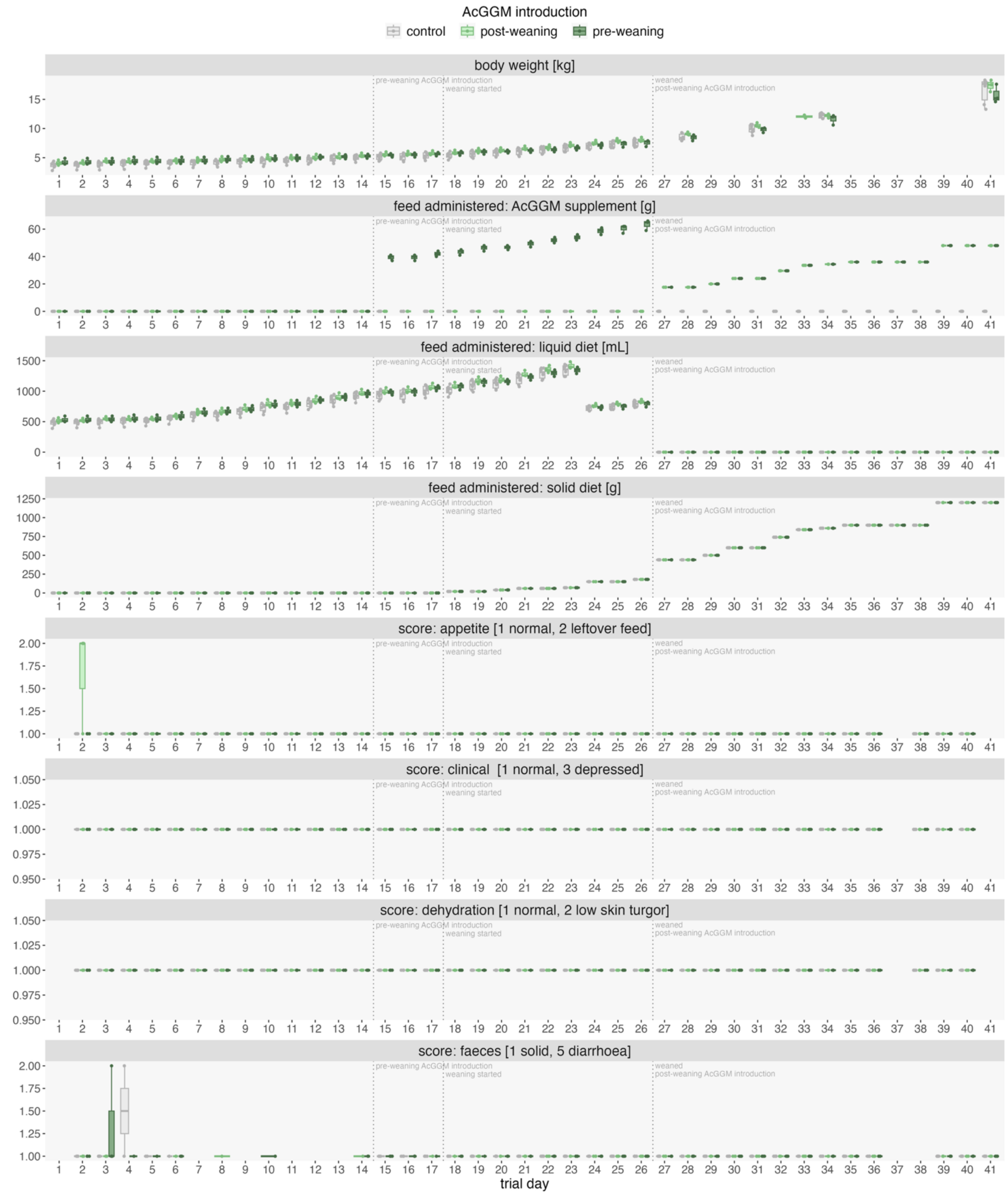

***Figure S3.1****. Animal performance and administered feed throughout the feeding trial.*

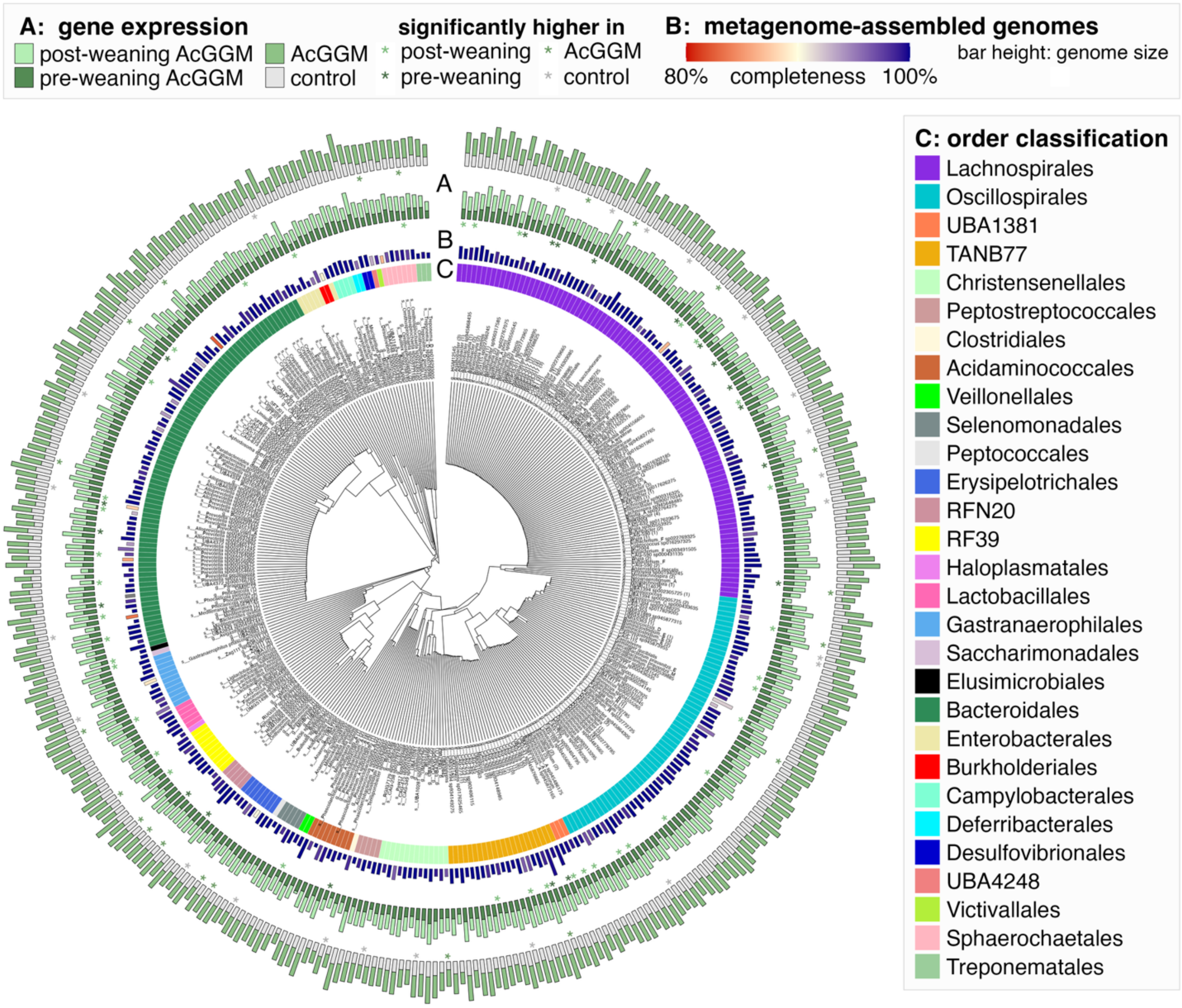

***Figure S3.2**. Phylogenetic tree of the 370 bacterial populations in the MAG catalogue. The most specific taxonomic classification is used as branch labels. Repeated taxon identifiers are numerated by order of appearance. **A)** Sum of metatranscriptomic reads mapped to each MAG in animals fed AcGGM, split into fibre introduction with or without AcGGM (outer ring) and before and after weaning (inner ring). Significant differences (LFC>1, FDR p-value<0.05) in activity levels are marked with asterisks below each bar, with colour indicating which of diet group had the most transcriptionally active population of the MAG. **B)** Genome sizes (bar height) and completeness (colour) of MAGs. **C)** Taxonomic classification of MAGs on the order level.*

The remainer of **Suppl. 3** contains overviews of differentially abundant features from metagenomic, metabolomic, and host transcriptomic layers. Due to the high number of differentially expressed microbial genes, metatranscriptomics is not included. Lists of these features are instead found on GitHub (jennymerkesvik/3domics_wp3-2).

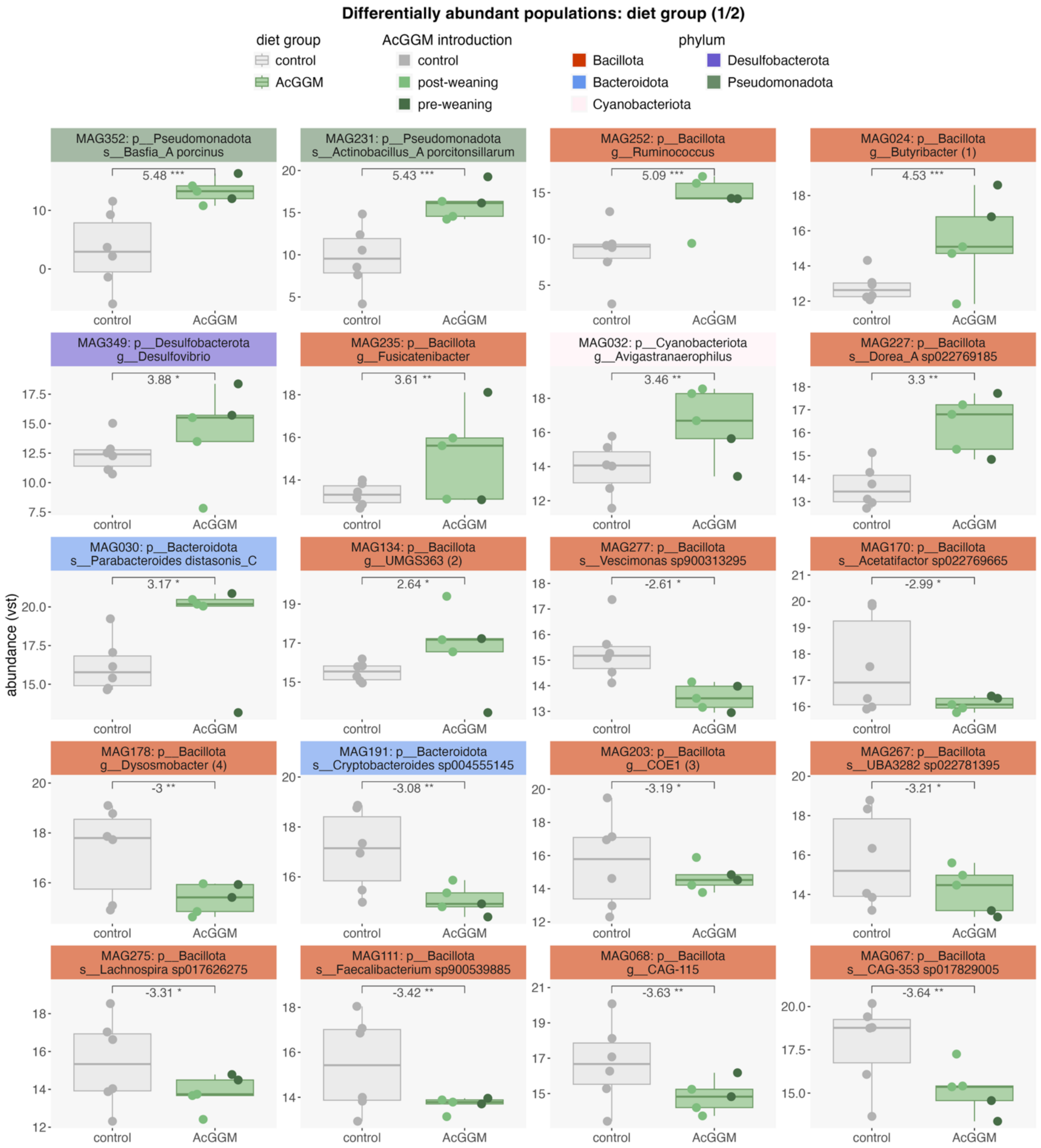

***Figure S3.3****. Different**ially abundant metagenome-assembled genomes in microbiomes of animals given different diets. Significance is indicated with horizontal bars, reporting log2 fold change and false discovery rate-adjusted p-values (* < 0.05, ** < 0.01, *** < 0.001). Continued on next page.*

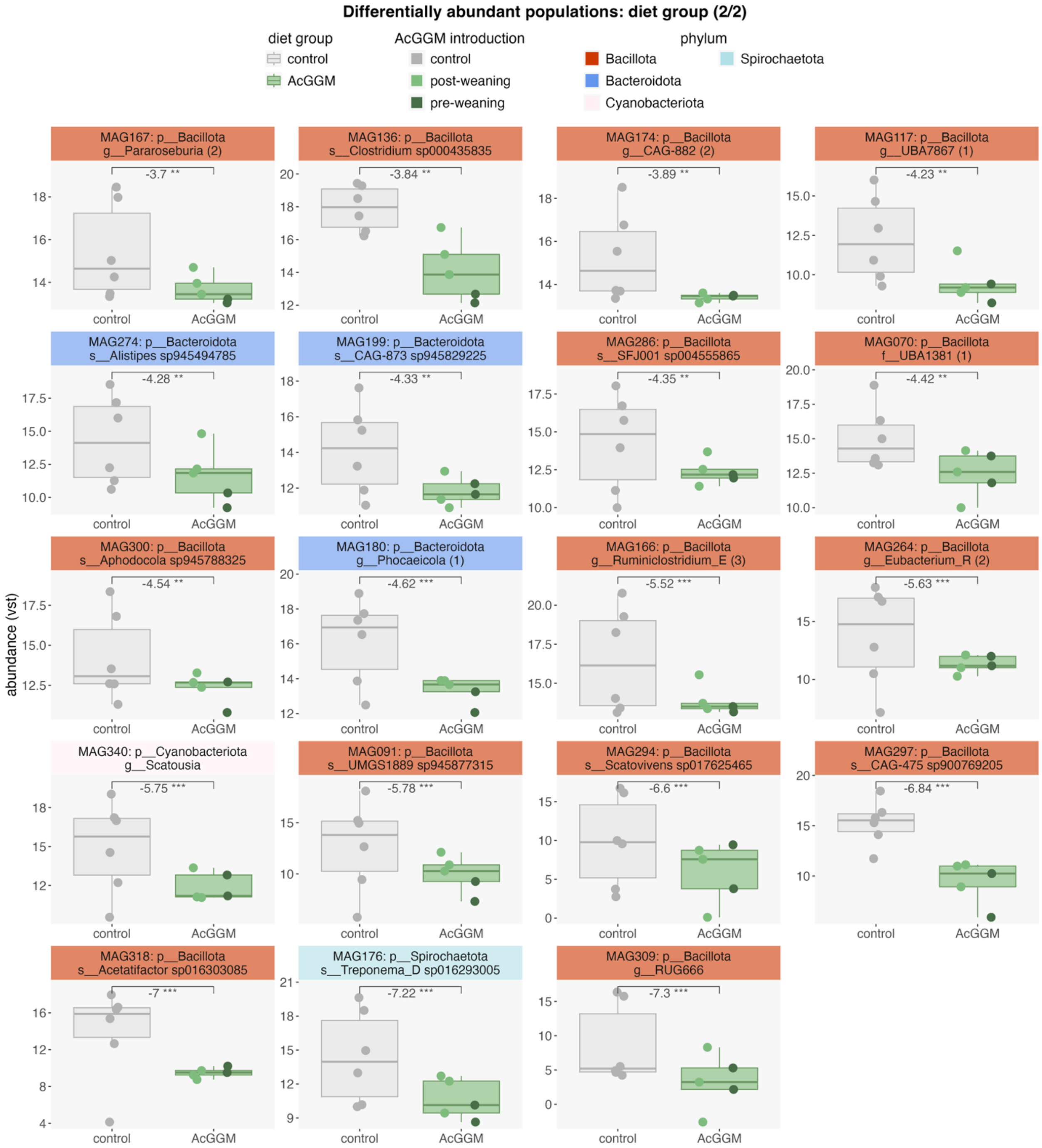

***Figure S3.3*** *continued (end).*

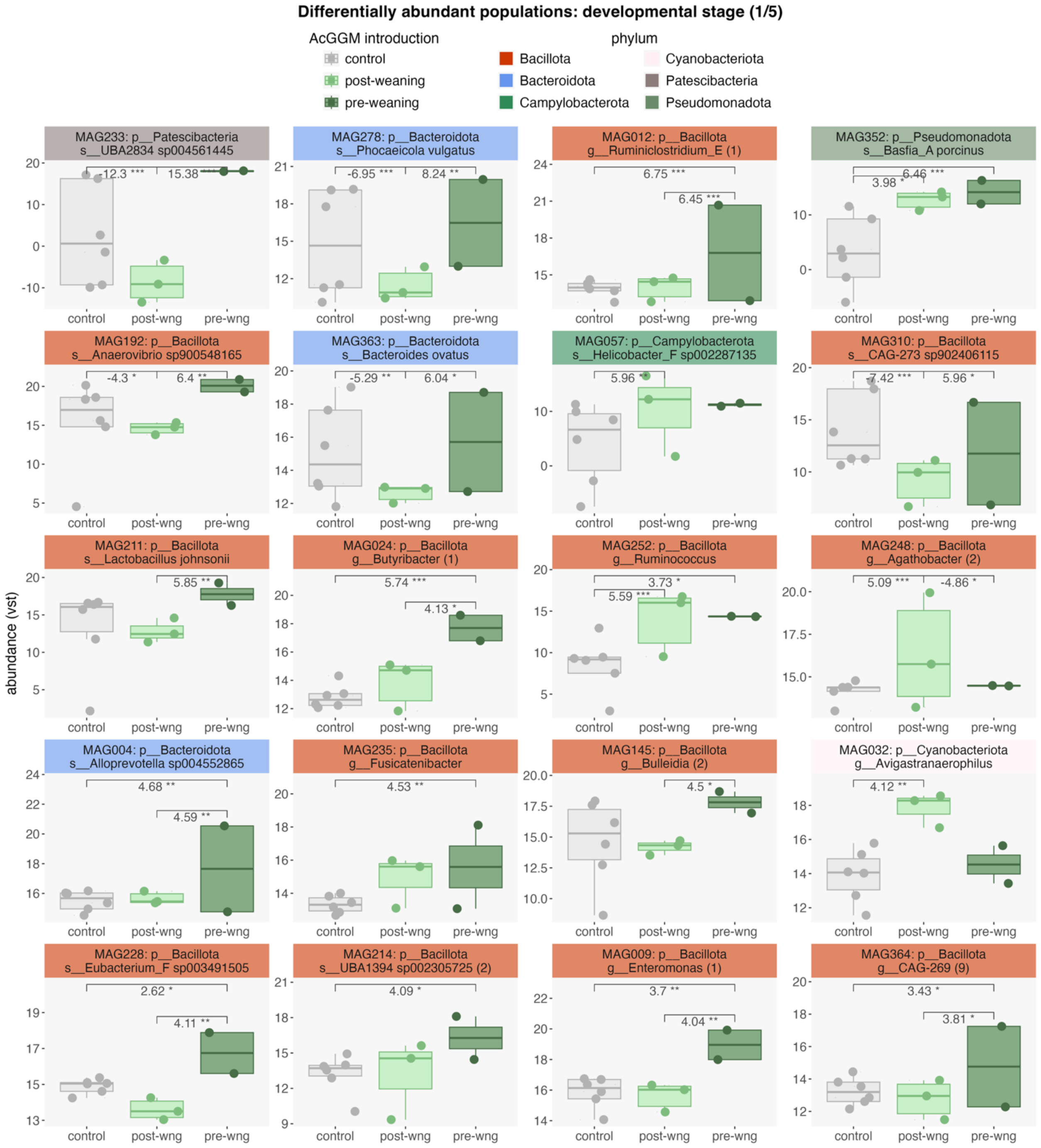

***Figure S3.4**. Differentially abundant metagenome-assembled genomes in microbiomes of animals introduced to acetylated galactoglucomannan fibres starting at different developmental stages. Significance is indicated with horizontal bars, reporting log2 fold change and false discovery rate-adjusted p-values (* < 0.05, ** < 0.01, *** < 0.001). Continued on next page.*

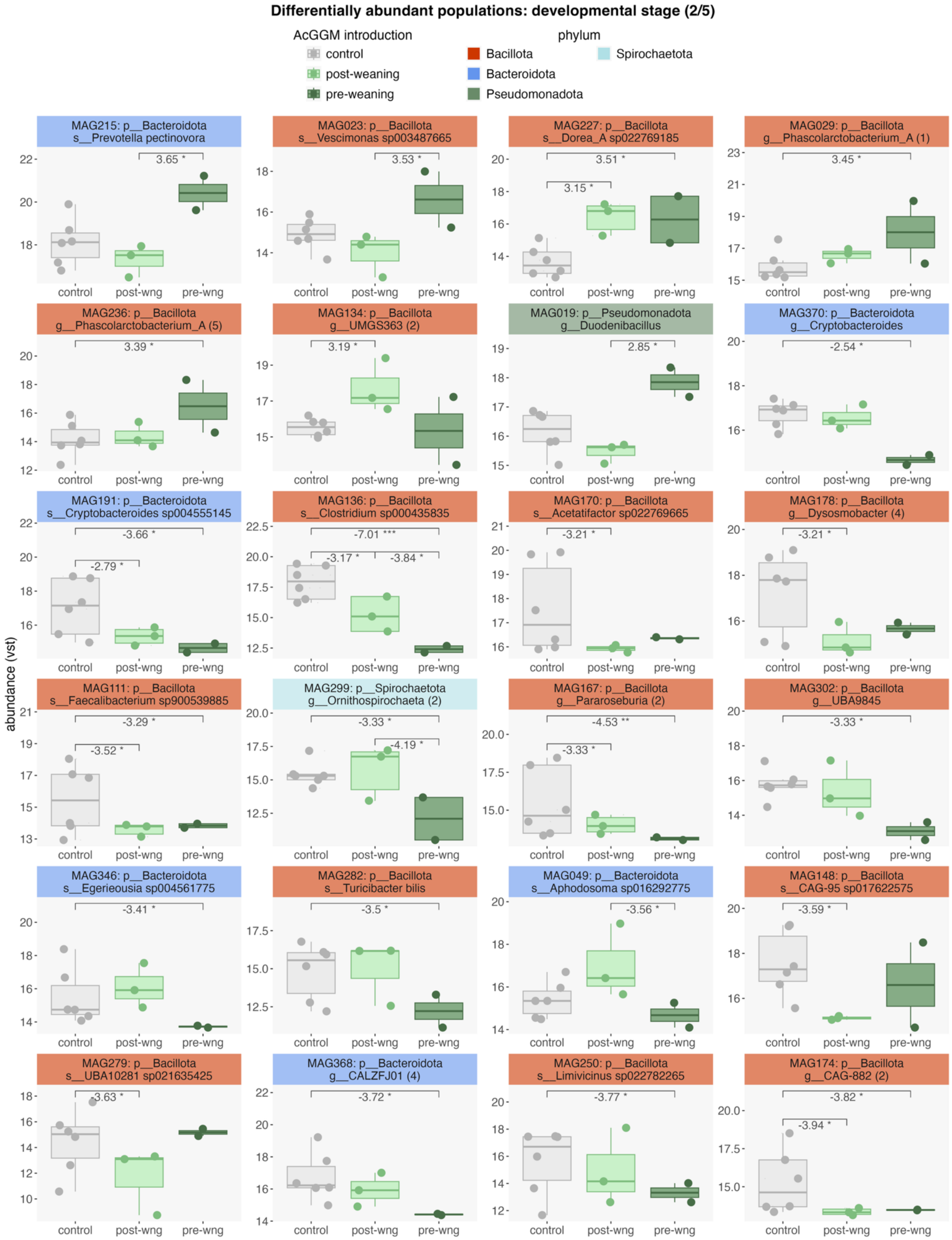

***Figure S3.4** continued.*

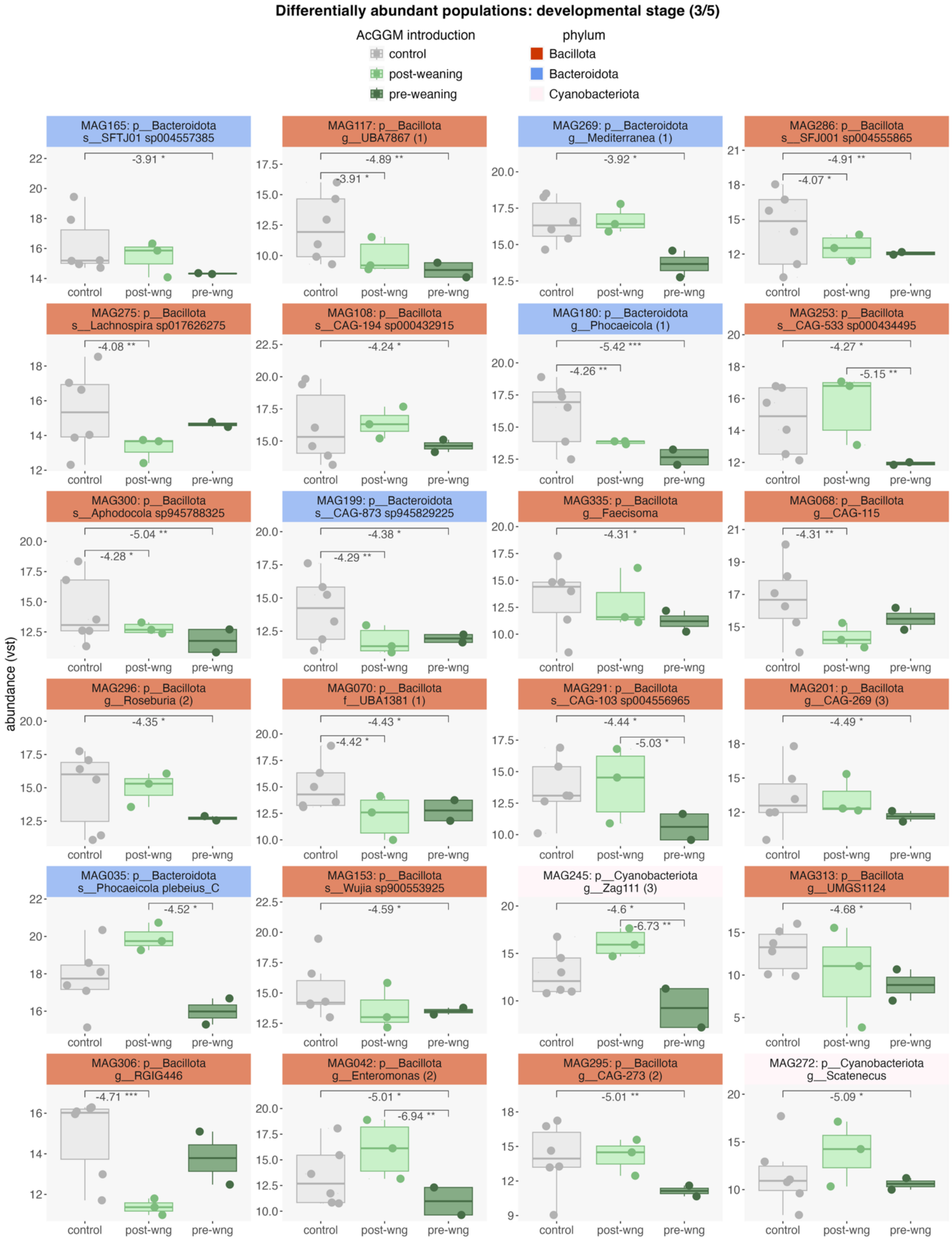

***Figure S3.4*** *continued.*

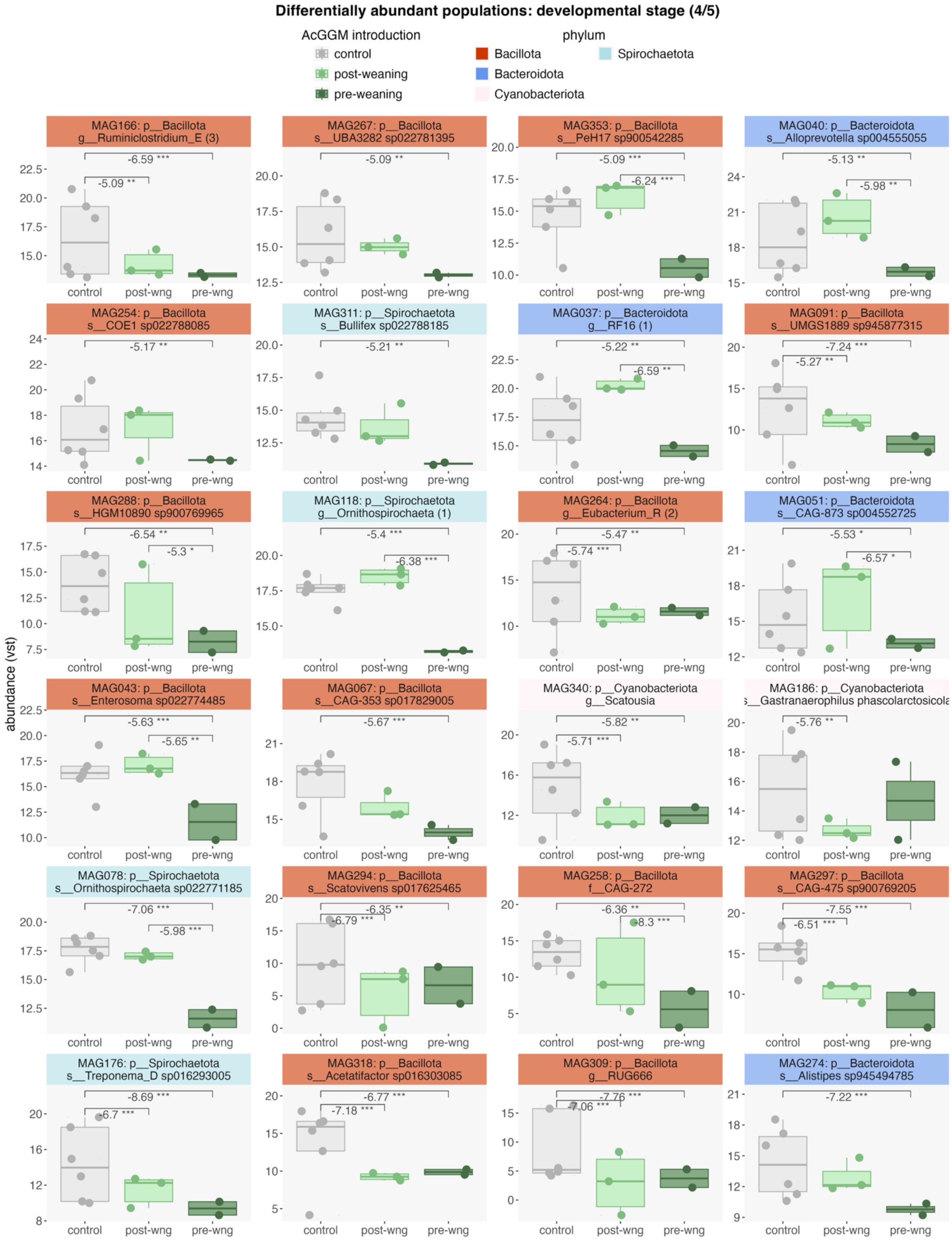

**Figure S3.4** continued.

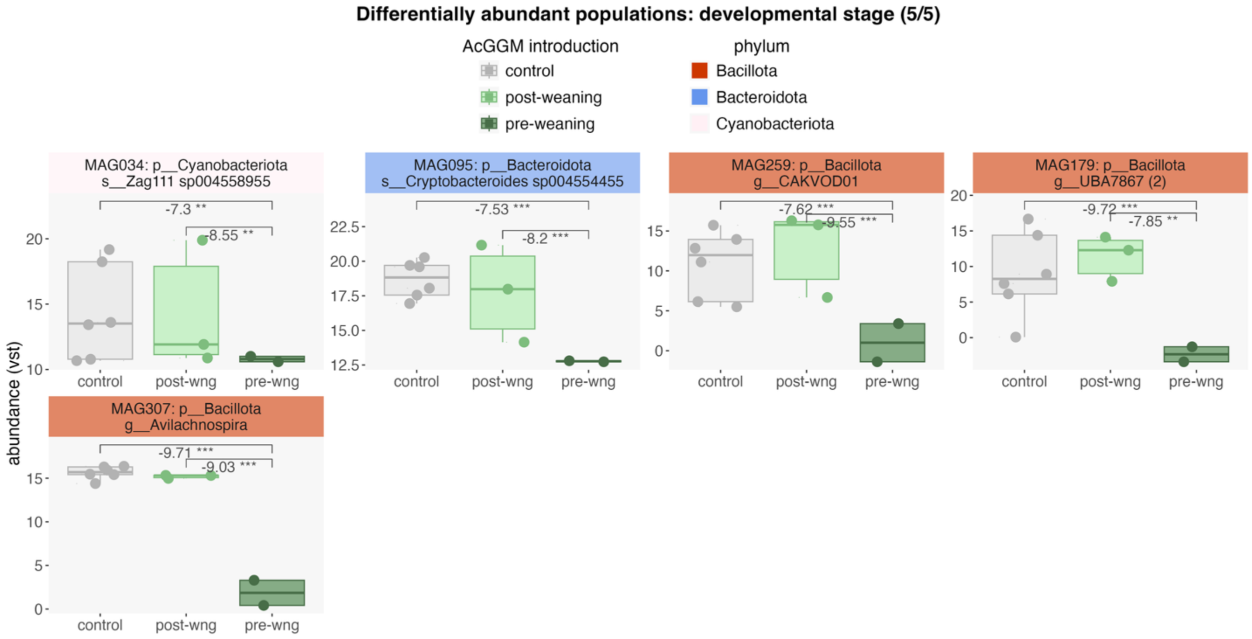

***Figure S3.4*** *continued (end).*

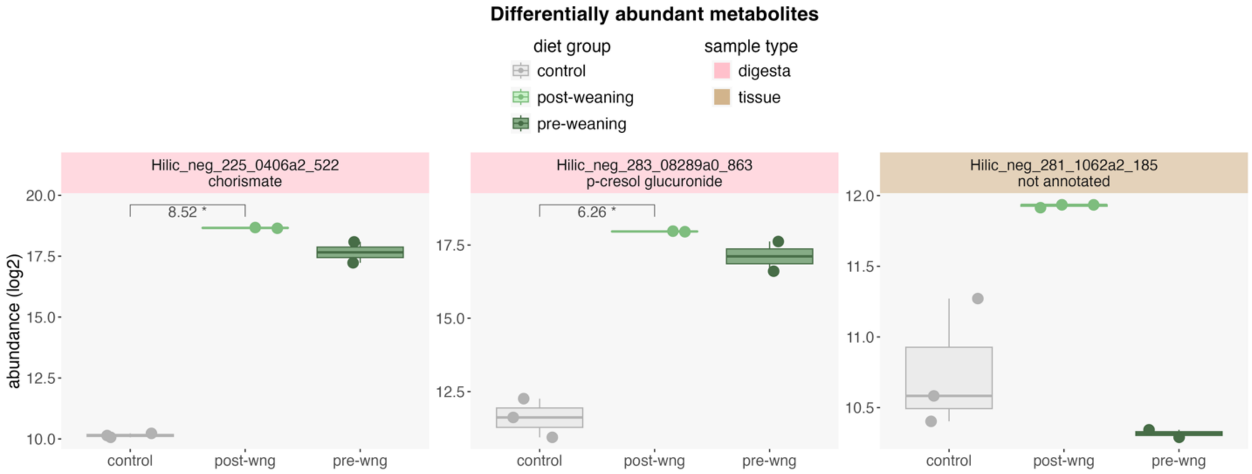

***Figure S3.5**. Differentially abundant metabolites in digesta or tissue samples of animals introduced to acetylated galactoglucomannan fibres at different developmental stages. Significance is indicated with horizontal bars, reporting log2 fold change and false discovery rate-adjusted p-values (* < 0.05, ** < 0.01, *** < 0.001).*

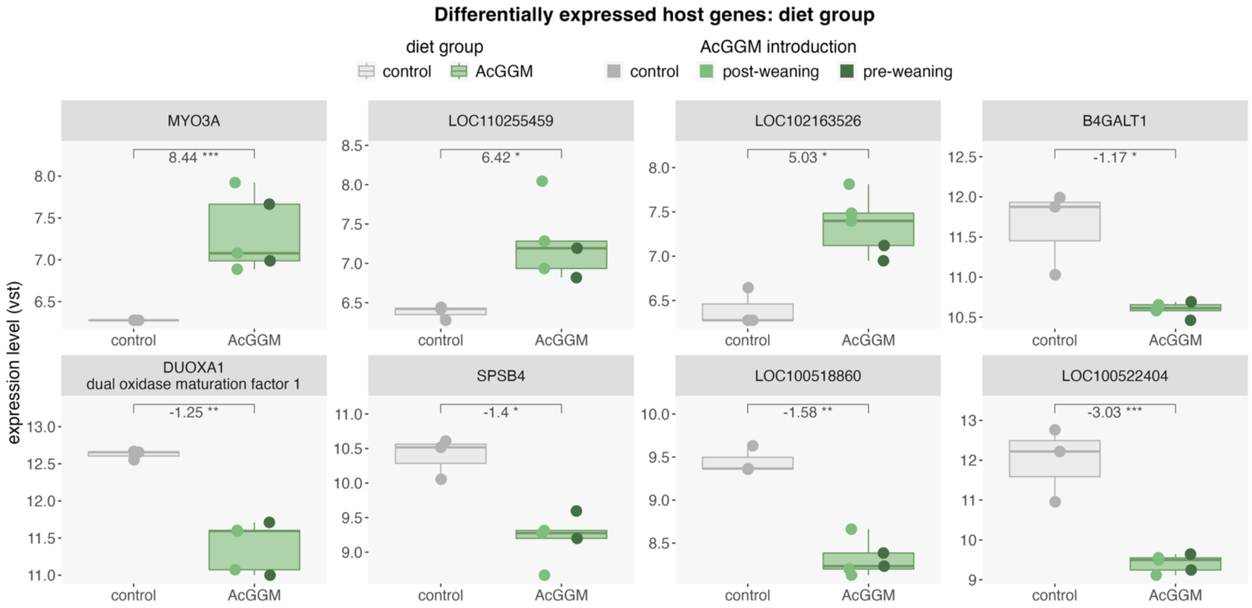

***Figure S3.6**. Differentially expressed genes in caecal gut wall samples of animals given different diets. Significance is indicated with horizontal bars, reporting log2 fold change and false discovery rate-adjusted p-values (* < 0.05, ** < 0.01, *** < 0.001).*

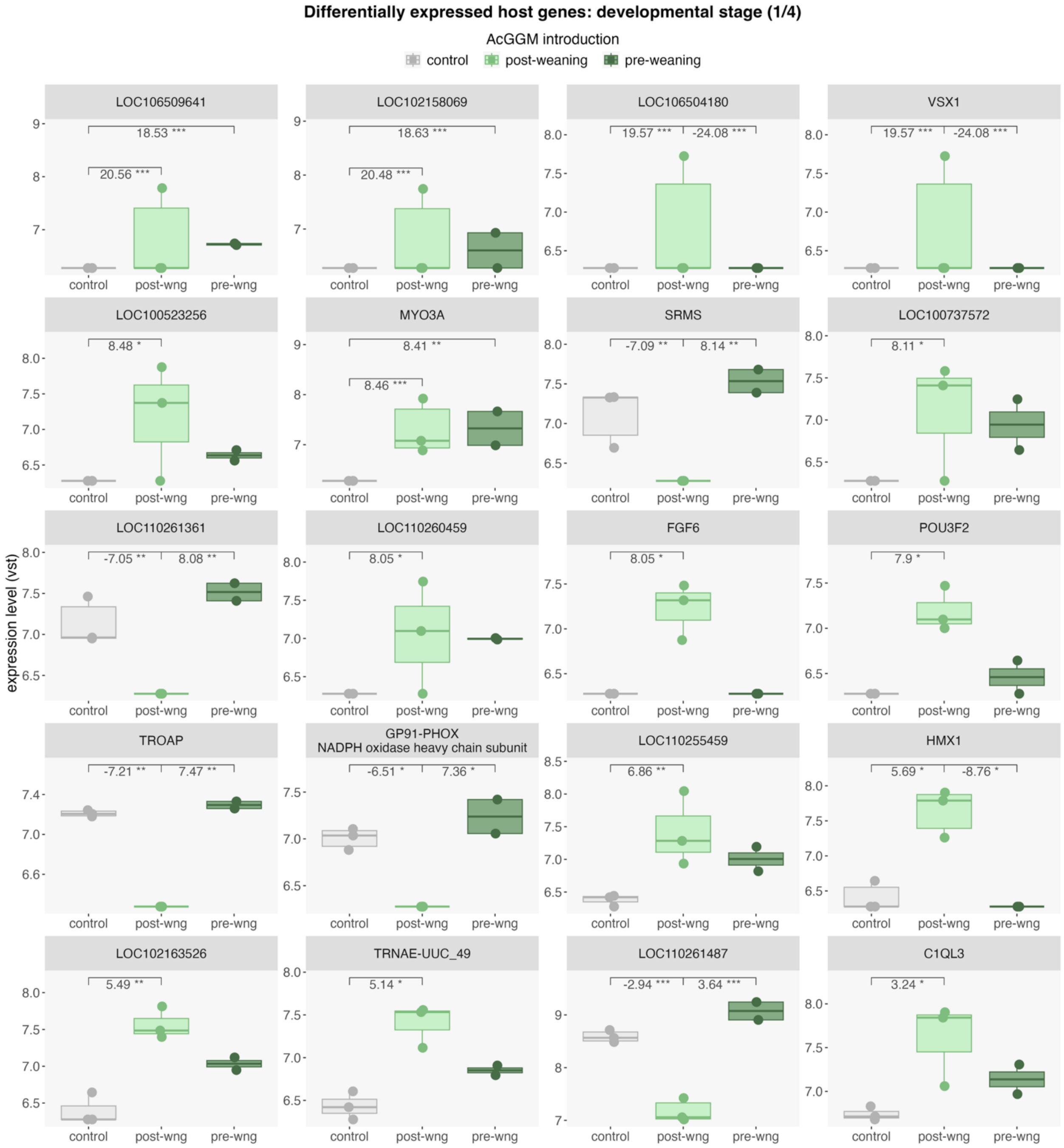

***Figure S3.7****. Differentially expressed genes in caecal gut wall samples of animals introduced to acetylated galactoglucomannan fibres starting at different developmental stages. Significance is indicated with horizontal bars, reporting log2 fold change and false discovery rate-adjusted p-values (* < 0.05, ** < 0.01, *** < 0.001). Continued on next page.*

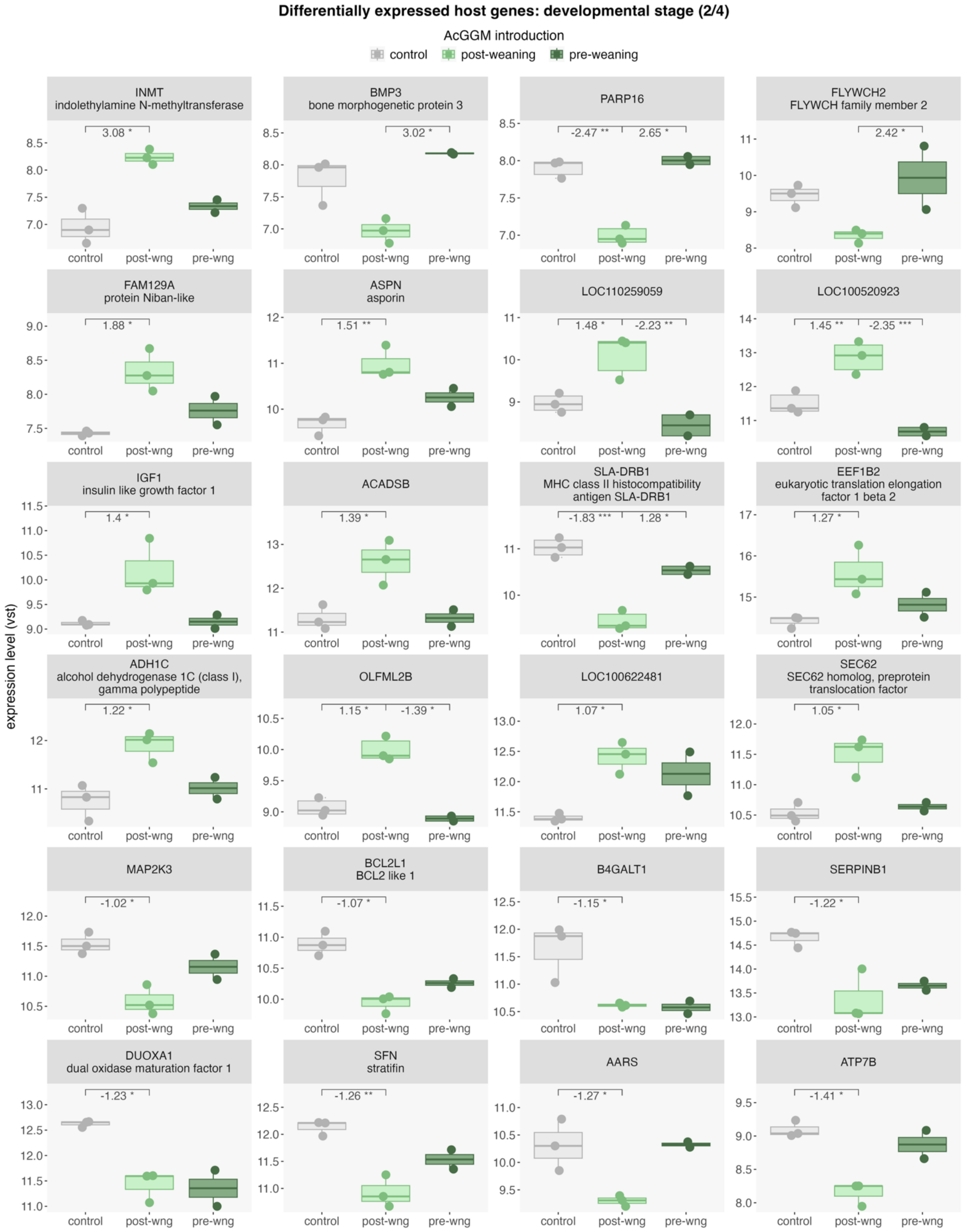

***Figure S3.7*** *continued.*

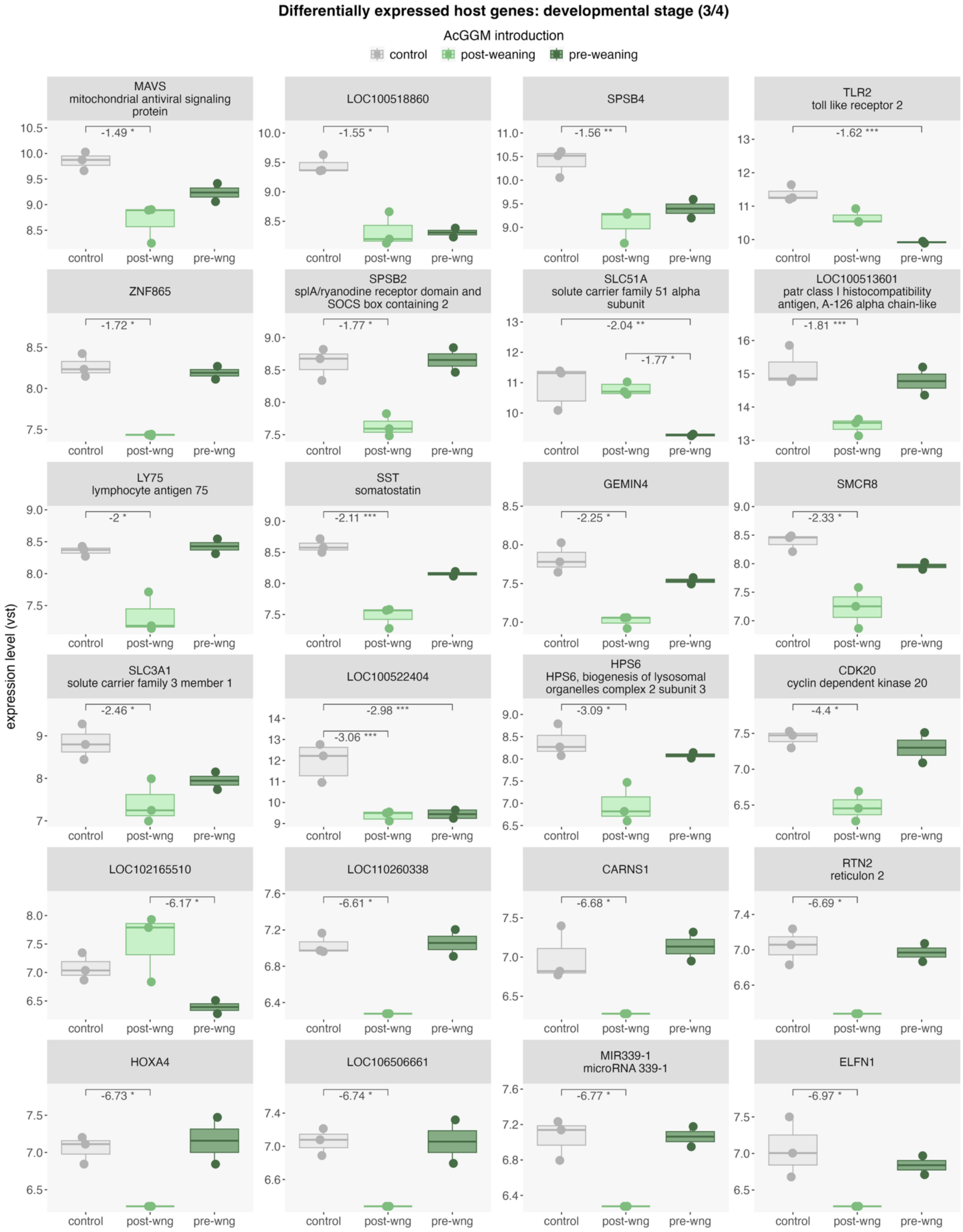

***Figure S3.7** continued.*

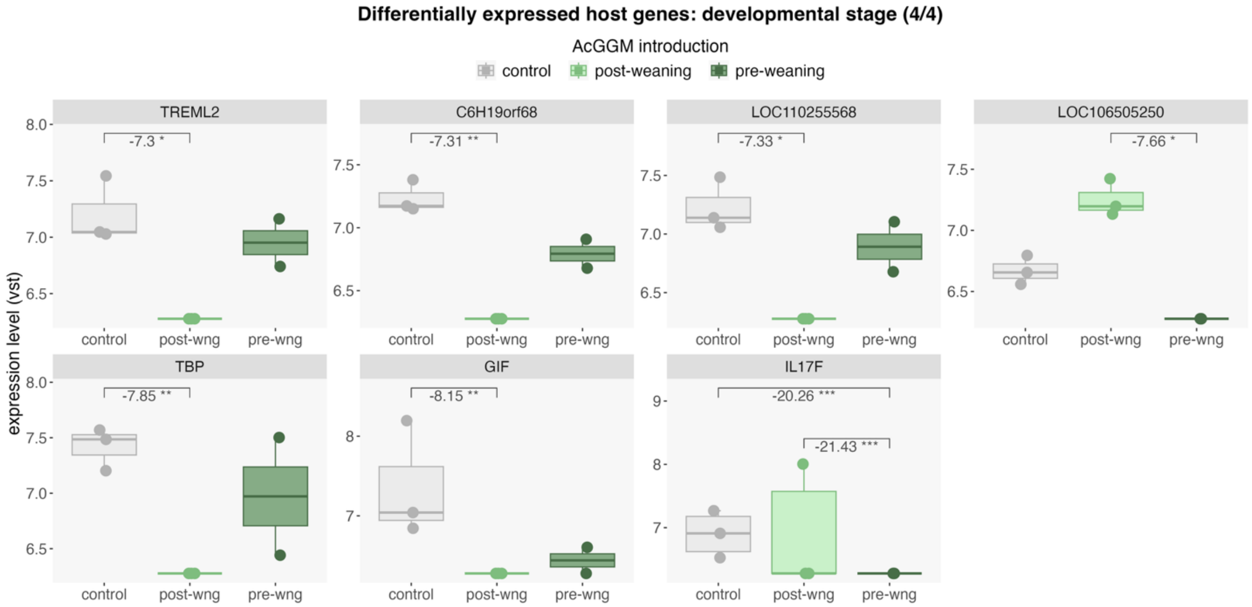

**_Figure S3.7_** _continued (end)._
